# Mitigating *Enterotoxigenic Escherichia coli* (ETEC) infection with Probiotic and Synbiotic Treatments: A murine and in vitro study

**DOI:** 10.1101/2025.08.19.671123

**Authors:** Sohini Sikdar, Paulami Dutta, Debmalya Mitra, Bidisha Pal, Hemanta Koley, Shanta Dutta

**Author notes:** **Correspondence:** Shanta Dutta, ICMR-National Institute for Research in Bacterial Infections (ICMR-NIRBI), P-33 C.I. T Road, Beleghata, Kolkata 700010, West Bengal, India.

## Abstract

**Background & Aims:** *Enterotoxigenic Escherichia coli* (ETEC) strain H10407 is a major causative agent of Diarrhoeal diseases. This study investigates the inhibitory effects of selected probiotics (*Lactobacillus casei*, *Lactococcus lactis*, and *Bifidobacterium bifidum*) and their synbiotic combinations with inulin on ETEC proliferation, adhesion, intestinal colonization, and host immune response.

**Methods:** ETEC H10407 was cultured in Luria-Bertani broth and co-cultured with probiotic strains to assess growth dynamics. Adhesion and competitive inhibition assays were performed using INT 407 intestinal epithelial cells. A murine infection model (C57BL/6) was employed to evaluate pathogen colonization, diarrheal severity, stool consistency, histopathological changes, and immune modulation. Synbiotic efficacy was further tested by combining probiotics with the prebiotic inulin. Additionally, fecal microbiota transfer (FMT) was conducted to examine microbiome-mediated protective effects.

**Results:** Probiotic co-culture significantly inhibited ETEC growth from 2 h onward without impairing probiotic viability. All probiotics markedly reduced ETEC adhesion to INT 407 small intestinal cells. In vivo, probiotics lowered ETEC colonization by 81.34%, while synbiotics conferred an additional 29.68% reduction. Synbiotics prevent watery stool and preserve ileal architecture, with reduced goblet cell loss and neutrophil infiltration. Synbiotics significantly (p<0.0001) decreased CD3+/CD4+ Th1 cells in spleen and mesenteric lymph nodes, reduced pro-inflammatory cytokines and chemokines, and restored tight junctional proteins.

**Conclusion:** Synbiotic supplementation offered superior protection against ETEC-induced diarrhea compared to probiotics or prebiotics alone, highlighting its potential as a non-antibiotic prophylactic strategy. These findings support the role of microbiome-targeted interventions within the One Health framework to address enteric infections and antimicrobial resistance.

## Introduction

*Enterotoxigenic Escherichia coli* (ETEC) is an enteropathogen, responsible for a significant cause of bacteria-associated diarrhoeal illness. According to the Centers for Disease Control and Prevention (CDC), Infection with ETEC is the major cause of traveler’s diarrhoea and among children in low-income countries. As per the World Health Organization (WHO), globally, childhood Diarrhoea is reported in 1.7 billion cases with an estimated 525000 deaths per year. According to UNICEF, most of the Diarrhoeal deaths among children are from South Asia and sub-Saharan Africa. Though ETEC infection is not limited to any age group, it poses a significant threat, particularly among children who are yet to be immunized, with prior infections (1). ETEC causing Traveler’s diarrhoea affects 10–40% of travelers (gender unbiased), with incidence depending on destination, country of origin, length of stay, and travel season (2), (3), (4). The genetics of people, though, greatly influence disease susceptibility (5) and multiple episodes can occur during one trip (6).

*Enterotoxigenic Escherichia coli* (ETEC) attaches to intestinal epithelial cells using fimbriae-associated proteins known as colonization factors (CFs), which include specific adhesins (7). This bacterial attachment initiates the release of toxins—either heat-labile (LT) or heat-stable (ST) enterotoxins. ETEC also secretes the protease EatA and the metalloprotease YghJ, which promote mucin degradation, thereby facilitating ETEC outer membrane vesicles (OMVs) access and internalization into the host’s lipid raft regions. The LT and ST toxins activate host adenylyl cyclase and guanylate cyclase, respectively, increasing intracellular cAMP and cGMP levels. These signaling cascades disrupt ion transport, leading to excessive secretion of electrolytes and water into the intestinal lumen, culminating in watery diarrhoea in travelers (8). Currently, there is no licensed vaccine available against ETEC infection (9), (10). The complex nature of ETEC and the lack of detailed knowledge about the molecular mechanism of pathogenesis pose many challenges that restrict the development of a vaccine that is broadly protective, inexpensive, and practical (11).

ETEC transmits its pathogenicity through two primary endotoxins: heat-stable (ST) and heat-labile (LT). ST is a peptide consisting of 19 amino acids that has a low immunogenicity and significant toxicity. LT comprises one LTA subunit and five LTB subunits and forms a macromolecule. Structurally and functionally LT holotoxin has a close resemblance to cholera toxin (CT). There are ETEC strains expressing either LT or ST or both (12), (13).

Moreover, indiscriminate and irregular exposure to antimicrobial agents has also resulted in antimicrobial resistance in ETEC. Therefore, it is necessary to reduce the burden of the disease with an alternative approach that would reduce the rate of antimicrobial resistance and should have some health benefits. The primary limitation in vaccine development against ETEC was the heterogeneity of adhesins (14), (15), (16), (17). The only oral vaccination that has been developed so far and is undergoing a phase 2b trial study is ETVAX (18), (19). Here in our study, we attempted to mitigate ETEC infection-induced pathologies using an alternative approach with probiotic bacterial strains and their combination with prebiotics, also referred to as synbiotics.

Probiotics are live beneficial microorganisms that are often used as an alternative treatment for many gastrointestinal diseases such as Inflammatory Bowel Disease (IBD) (20), (21), (22), colorectal cancer (23), (24), antibiotic-associated diarrhoea (25), (26) and enteric infections (27), (28). It has been observed that probiotics reduce microbial toxicity in the intestine directly or indirectly via bacteriocins, and improve our gut functions and our innate immunity (29). Strains of probiotic species are known to increase anti-inflammatory cytokines, like IL-10 and TGF-β, and decrease pro-inflammatory cytokines such as IL-6, IL-8, and TNF-α (30), (31), (32), (33). Probiotic strain has been shown to increase tight junction protein like ZO-1 and transmembrane protein like occludin in duodenal samples, creating a plausible approach to use probiotics to enhance gut epithelial barrier in humans (29). On the other hand, prebiotics are “non-digestible food ingredients that are beneficial to the host by selectively stimulating the growth or activity of one or a limited number of bacteria residing in the colon.” (34). Prebiotics require a colony of potentially beneficial microbes to ferment them specifically (35). Fermentation of prebiotics produces short-chain fatty acids (SCFAs) like lactic acid, butyrate, and propionic acid, which help to decrease colon pH and positively influence the intestinal epithelial barrier (36), (37), (38). Fructo-oligosaccharides (FOS) & Galacto-oligosaccharides (GOS) are the most commonly used prebiotics, which can be fermented by strains of *Bifidobacterium*, *Lactobacilli,* and *Aspergillus* species (39). This study used Inulin, which is a plant-based fructo-oligosaccharide with a low caloric value (1.5kcal/g or 6.3kJ/g) and non-digestibility. Inulin in the presence of intestinal bacteria transforms into short-chain fatty acids (like acetate, propionate, and butyrate), lactate, bacterial fuel and gases (40), (41). SCFAs, therefore, are consumed by Bacteria and partially used up by the host cells (42). According to studies, probiotics and prebiotics work better together to shorten the duration of Diarrhoea and hospitalization episodes. When taken orally, synbiotics produce an inhibitory substance, lower the risk of infection, and improve the hosts’ nutritional status. Different prebiotics are used in combination with a different probiotic to form synbiotics i.e. The first three are *Lactobacillus plantarum* 299 with oat fiber; the second is *Lactobacillus sporogens* combined with fructo-oligosaccharides; the third is Synbiotic 2000, which has inulin, betaglucans, pectin, and resistant amide along with each of *Pediococcus*, *Leuconostoc*, *Lactobacillus species*, fourth, *Pediococcus pentoseceus*, *Leuconostoc mesenteroides*, *Lactobacillus species*, and 2.5 g of inulin, oat fiber, pectin, and resistant starch; Fifth, inulin (SYN1) + Oligofructosa + *Lactobacillus rhamnosus* GG and *Bifidobacterium lactis* Bb12; vi) Golden *Bifid*: *Bifidobacterium bifidum*, *Lactobacillus bulgaricus*, and *Streptococcus thermophilus* with FOS (43). Human intestinal probiotic bacteria *B. fragilis*, *S. parasanguinis* showed greater growth in Inulin than in media supplemented with glucose only. On the contrary, *E. coli* can grow significantly faster in glucose medium but not in inulin-supplemented medium (44), (45), (46), (47), (48), (49). Synbiotics are known to interfere with the adhesion and attachment of pathogens to gut mucosa, thereby preventing disease and gut dysbiosis (50). Adhesion in the intestinal epithelial cells is the key step for colonization, which leads to the disruption of the gut barrier and integrity and causes intestinal infection (50).

Therefore, this study assessed the efficacy of different Synbiotics in reducing the severity of ETEC infection in mice.

## Material & Methods

### Chemicals and reagents

De Man, Rogosa, and Sharp (MRS) broth (BD 288130) and agar (M641, Himedia), Luria Bertini broth (BD 244620), Bifidobacterium broth (M1395, Himedia), and MacConkey agar (BD 211387) were purchased from BD Biosciences. DMEM cell culture medium was purchased from Sigma-Aldrich. Fetal Bovine Serum (FBS) was from Gibco. ELISA kits for mouse TNF-α (BD OptEIA 555268), IL-6 (BD OptEIA 555240), IFN-γ (BD OptEIA 555138), CXCL-1 (R&D Systems DY453-05), and IL-10 (BD OptEIA 555252) were purchased from BD Biosciences. IHC and FACS Antibodies FITC-tagged anti-CD3 (cat #100204, BioLegend), PerCP-tagged anti-CD4 (cat#100537, BioLegend), and PE-tagged anti-CD8a (cat#100707, BioLegend) were purchased from BD Biosciences.

### Enterotoxigenic *Escherichia coli* and probiotic bacterial strains

The probiotic strains used in this study were *Lactobacillus casei* ATCC 393, *Lactococcus lactis* ATCC 49032, and *Bifidobacterium bifidum* ATCC 11863. Watery diarrhoea was induced with the ETEC H10407 strain, which is both LT and ST positive.

### Growth inhibition of ETEC in the co-culture system with the probiotics

To investigate the effect of probiotic strains on the growth kinetics of ETEC H10407, we performed a co-culture assay. For this experiment, bacterial pure cultures and co-cultures were prepared. For pure culture, 50 ml of single-strength Tryptic Soy broth (TSB; BD, USA) and 50 ml of single-strength De Man-Rogosa-Sharpe (MRS; HiMedia) broth were inoculated with 1×10^8^ CFU of ETEC and *Lactobacillus casei*, respectively, and were incubated at 37℃ with a constant shaking of 150 rpm. Similarly, for a 50 ml co-culture medium, 25 ml of each of double-strength TSB and MRS broth (TSB: MRS-1:1) was mixed and inoculated with 1×10^8^ CFU of both ETEC and *Lactobacillus casei* together. The whole procedure was repeated identically for the *Lactococcus lactis* and *Bifidobacterium bifidum* strains. 100 µl of medium from each pure and co-cultures was collected at 0, 2, 4, 8, 10, 12, 24 h and lawn cultured in appropriate agar plates to determine the viable colony count. Plates were incubated overnight at 37℃ for colonies to grow. The tests were performed in triplicate.

### Cell culture

INT 407 (originally a human embryonic small intestinal cell line) cells were cultured in DMEM supplemented with 10% FBS and kept in a 5% CO2 incubator at 37°C. The cell culture media was changed at regular intervals. Confluent cells were trypsinized before experiments and seeded in a certain number with the help of a haemocytometer.

### Adhesion assay

For the ETEC adhesion assay, the INT 407 cells were seeded at a density of 10^6^ cells/well. The day before the experiment, the cell culture medium was replaced with antibiotic-free medium. From each fresh batch of the bacterial cultures, ETEC H10407, *L. casei*, *L. lactis,* and *B. bifidum* were harvested by centrifugation at 10,000 rpm for 8 min at 4°C and introduced to cultured cells at MOI 20. In this assay, we assessed the adhesion inhibition and competitive inhibition of ETEC H10407 by different probiotic strains. For the adhesion inhibition assay, probiotics were introduced 2 hours before ETEC addition to the cells, and for the competitive inhibition assay, the probiotics and ETEC were introduced at the same time to the cells (51). After the ETEC introduction, cells were incubated for 2 hours. Cells were then washed 5 times with cold 1X PBS buffer. Cells were lysed using Triton-X-100 and spread on. The viable colonies from each cultured organism were counted from plates of serial diluents on MRS and Luria agar and used accordingly in the assay.

### Animal maintenance

We bought C57BL/6 mice from Kalyani in West Bengal, India. The mice were kept in a 12-hour light-dark cycle with an ideal temperature of 24° C, fed a regular chow diet, and given unlimited access to autoclaved drinking water. (Details of animal ethics No. NICED/CPCSEA/68/GO/(25/294)/2022-IAEC/SD/1).

### Experimental design

Four to six weeks old Balb/c mice were grouped (n ≥ 5) into-(a) PBS control; (b) ETEC; (c) IN; (d) *L. casei*; (e) *L. lactis*; (f) *L. casei* + IN; (g) *L. lactis* + IN and (h) *B. bifidum* + IN. Mice were fed a standard chow diet ad libitum and autoclaved drinking water during the experimental period and reared in a 12-hour light-dark cycle condition and temperature (25° C ± 2). Mice in IN, *L. casei* + IN, and *L. lactis* + IN, and *B. bifidum* + IN groups were fed Inulin (2.5g/ kg body wt.) from day 0 to day 14. The dose of inulin was referred by various authors (52), (53). Mice in *L. casei*, *L. lactis*, *B. bifidum*, *L. casei* + IN, and *L. lactis* + IN and *B. bifidum* + IN were orally administered with 10^8^ CFU/ml of respective probiotic strains from day 7 to day 14. All the groups except for the control were infected with 10^9^ CFU/ml of ETEC strains on day 14.

### ETEC colonisation in the ileum

All mice were sacrificed after 20 hours of infection. Portions of the ileum from mice from all the groups were taken and homogenized in PBS and plated onto the MacConkey agar plate. Colonies were further validated by real-time PCR using specific primers against both the ST and LT genes of the ETEC H10407 strain (data not shown).

### Histological analysis of ileal sections

The Small pieces of ileum were first cleaned with phosphate-buffered saline (PBS), fixed with 10% formaldehyde solution, and then left for 48 hours to examine the degree of intestinal inflammation in the various groups. The tissues were embedded in paraffin after being dehydrated in alcohol. Sections 5–6 µm thick were cut with an ultra-microtome and stained with hematoxylin and eosin dyes, which were then examined under a microscope. The Erben et al. (2014) method was used to determine the histopathological scores (54).

### Flow cytometric analysis of the T cell population

Mesenteric lymph nodes (MLN) and spleens were collected from each individual and homogenized in PBS. Both spleen and MLN lysates were then passed through 70µm cell strainers to obtain a cell suspension devoid of any debris. For MLN, the cell suspensions were centrifuged at 2000 rpm for 5 min to obtain the cell pellets. The cell pellets were then resuspended in FACS buffer (PBS with 10% FBS). On the other hand, Splenic filtrates were centrifuged along with Ficoll Paque Plus (Cytiva 17-1440-02) at a ratio of 1:3 for density gradient centrifugation at 2000 rpm for 30 min at zero breaks. The white cloudy phase of PMNs was then collected, leaving both the aqueous and ficoll layers. Further centrifugation was done to obtain the PMN pellets, and resuspended in FACS buffer. Cell suspensions were then divided into batches to stain with respective FACS antibodies, which were FITC-conjugated anti-CD3, PerCP-conjugated anti-CD4, and PE-conjugated anti-CD8 antibodies. The cell population was analyzed using FACS (BD Aria II).

### Enzyme-linked immune sorbent assay for cytokine profiling

Small portions of the ileum were homogenized to assess the inflammatory cytokines and chemokine levels. Tissues were homogenized in PBS 1X. Homogenates were centrifuged at 10000g for 1 min to collect the supernatants devoid of any tissue debris. The total concentration of the lysates was quantified using the BCA Protein Assay Kit (Thermo Scientific™ Pierce™, USA), and 50 µg lysate was used to perform ELISA for TNF-α, IL-6, IFN-γ, CXCL-1, and IL-10.

### Real-time PCR assay for Tight junctional gene analysis in the ileum

Total RNA was extracted from ileal sections using Trizol and quantified using a nanodrop machine (NanoDrop 8000 Spectrophotometer, ThermoFisher Scientific, USA). From 0.5 μg of RNA and reverse transcription master mix from the TaKaRa PrimeScript, the cDNA was prepared following the kit protocol. Real-time PCR was performed using specific primer sets for GAPD, ZO-1, Occludin, Claudin-2, and JAM-A.

Primer sets-

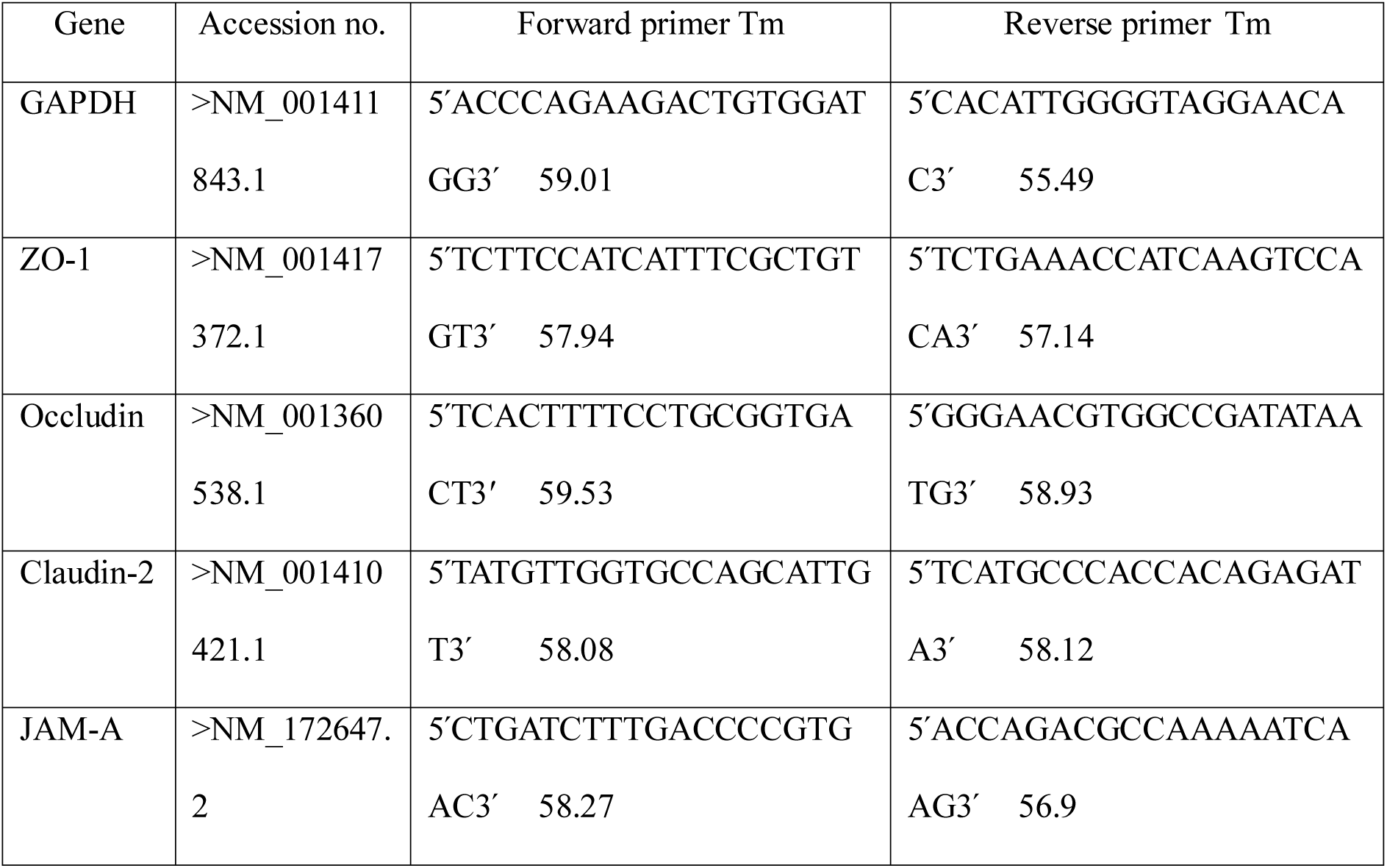

### Immunofluorescence assay for tight junctional proteins in the ileal sections

Paraffin-embedded colon tissue sections were first deparaffinized with Xylene and then rehydrated with graded ethanol. 0.5% Triton X-100 and 0.1% Sodium citrate solution were used to fix and permeabilize the tissue sections. Tissues were incubated for 1 hour in 5% sera in TBST as a blocking reagent. After that, tissue sections were incubated with primary antibody diluted with blocking buffer (1:200) overnight at 4 °C. Sections were washed three times with TBST buffer and incubated for 2 hours with fluorescent-tagged secondary antibody (1:200). DAPI (1:200) was used as the nuclear stain. Slides were finally mounted with glycerol and tissue sections were imaged using apotome fluorescence imaging system at 20X magnification.

### Fecal Microbiome Transplant (FMT) and pathogen clearance

Balb/c mice (n=3) were divided into three groups, and each group received a unique probiotic strain (*L. casei*, *L. lactis,* and *B. bifidum* separately) at a concentration of 10^8^ CFU/ml for 7 consecutive days. On day 6, another three sets of Balb/c mice (n=3) were infected with ETEC H10407 (10^9^ CFU/ml). On day 7, from each probiotic-fed mouse group, the fecal pellets were kept separately. Fecal pellets from each group were homogenized in a sterile PBS buffer to prepare microbial soup and labelled as *L. casei* soup, *L. lactis* soup, and *B. bifidum* soup. On day 7, each set of ETEC-infected mice group received any of the three fecal microbial soups, thus labelled as *L. casei* FMT, *L. lactis* FMT, and *B. bifidum* FMT. From day 8 to day 17, the clearance of ETEC infection was monitored daily by assessing the count of ETEC H10407 in the fecal matter of each FMT-grouped mouse. Fecal pellets were collected and weighed and homogenized and diluted in PBS, and then spread onto the MacConkey agar plate for counting. **Statistical analysis:**

Experiments were conducted in triplicate, and data are expressed as mean ± standard error of the mean, one-way analysis of variance (ANOVA) was conducted, followed by Tukey’s multiple comparison tests using Graph Pad Prism 6 software (Graph Pad Software Inc., San Deigo, CA, USA) and difference with P < 0.05 was considered significant.

## Results

### Probiotics inhibit ETEC growth and survivability, and adhesion to the host cells

The growth of ETEC H10407 in Luria Bertani broth shows an exponential time-dependent growth until it was co-cultured with *L. casei*, *L. lactis* and *B. bifidum,* respectively. A regression of the survivability of ETEC in the growth medium supplemented with probiotic strains was observed 2 hours after probiotic inoculation and onward. Although the proliferation of probiotic strains-*L. casei*, *L. lactis* and *B. bifidum* shows little to no inhibition of growth and proliferation patterns in the co-culture setup compared to their pure cultures (Figure 1A-C). INT 407 small intestinal cells were co-cultured with all three probiotic strains separately. Cells received probiotics 2 hours prior to infection for the adhesion inhibition assay, and cells that received probiotics along with ETEC were used for the competitive inhibition assay. The adhered ETEC CFU data, acquired from the infection control groups, was taken as 100 percent, and the rest was calculated with respect to that. All three probiotics significantly (p<0.0001) inhibited ETEC adhesion to INT 407 cells when compared to the infection control group. In the adhesion inhibition assay, the percentage of adhered bacteria to the cells was reduced to 15.7% with *L. casei*, 12.26% with *L. lactis*, and 14.63% with *B. bifidum*. Similarly, in the competitive inhibition assay, the ETEC adhesion to the INT 407 cells was observed to be 14.28% with *L. casei*, 23.3% with *L. lactis*, and 22.92% with *B. bifidum* (Figure 1D-F).

**Figure 1:**
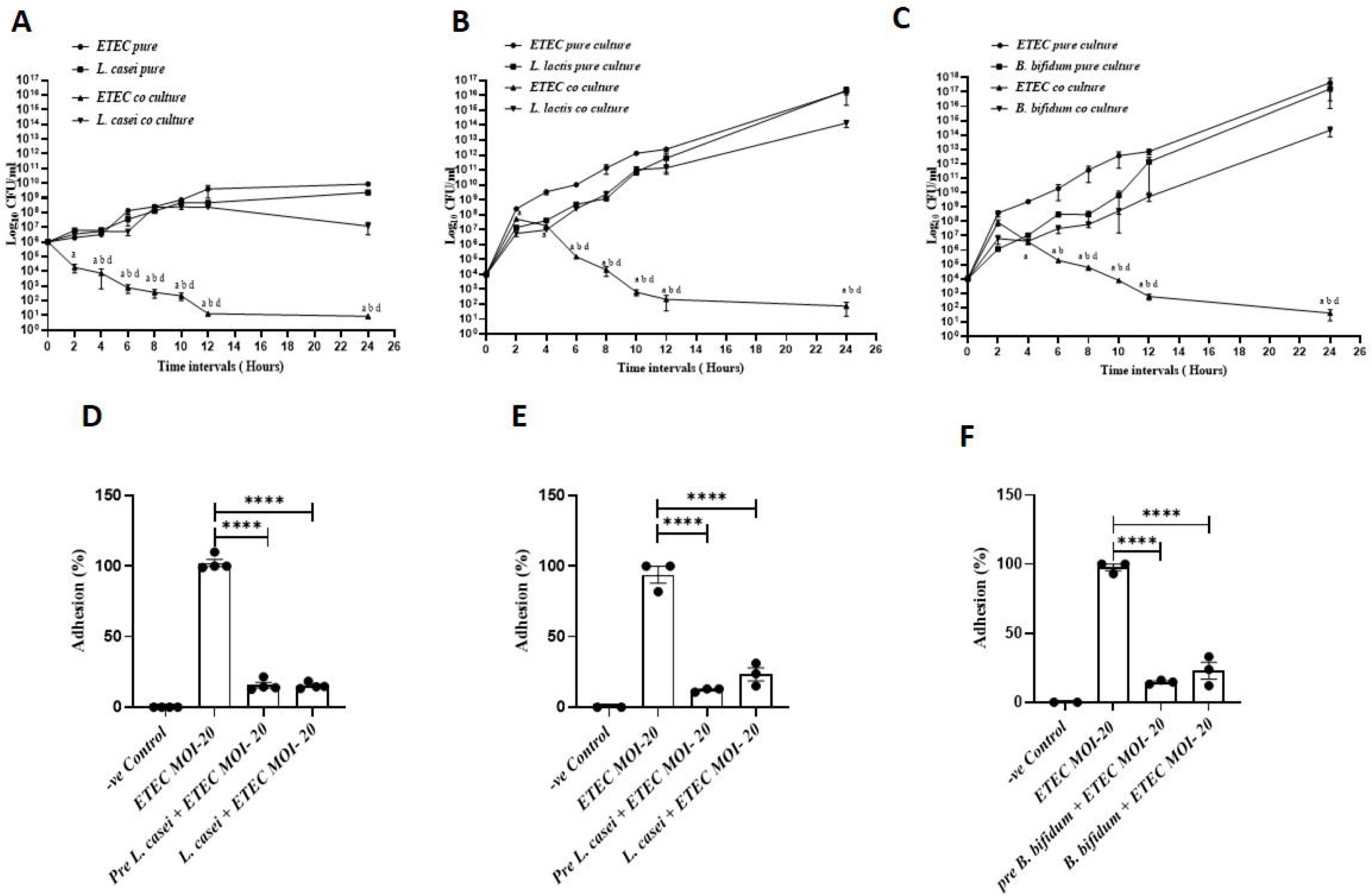
(A-C) Growth and survivability of ETEC and Probiotic strains *L. casei*, *L. lactis*, and *B. bifidum* in pure culture vs co-culture system. Bacterial strains were cultured and examined for 24 hours. A time-dependent growth pattern of bacterial strains was plotted on a Log^10^ scale. (D-F) ETEC H10407 adhesion to INT 407 small intestinal cells in the presence of *L. casei*, *L. lactis*, and *B. bifidum* (MOI 20), respectively. Pre L. *casei*, Pre *L. lactis*, and Pre *B. bifidum* terms were used in the adhesion inhibition assay. L. *casei*, *L. lactis*, and *B. bifidum* terms were used in the competitive inhibition assay. Adhesion Experiments were conducted thrice; the data are represented as Mean ± SEM.

### Synbiotics reduce ETEC H10407 load in the small intestine

In C57BL/6 mice, Diarrhoea was induced by the ETEC H10407 strain with a 10^9^ CFU/ml concentration per animal. After 20 hours of infection, the number of bacterial colonies detected on the MacConkey agar plate was significantly (p <0.0001) higher in the ETEC group compared to the control. RT-PCR and agarose gel electrophoresis further confirmed colonies. IN, *L. casei, L. lactis,* and *B. bifidum* probiotic groups showed a significant (p<0.0001) decrease in bacterial colonization with a percentage of 55.14433%, 71.37%, 81.34% and 67.60% when compared with the ETEC group. Also, the Inulin supplemented probiotic groups showed a significantly (p< 0.0001) lesser number of colonies compared to the ETEC positive group and even a further decrease (p< 0.004) in the bacterial colonization of 27.57%, 17.03% and 29.68% was noticed in the *L. casei* + IN, *L. lactis* + IN and *B. bifidum* + IN when compared to their corresponding probiotic group *L. casei, L. lactis,* and *B. bifidum,* respectively (Figure 2A). ETEC-infected mice displayed marked fluid accumulation in the intestinal lumen, which was evident from the presence of watery, yellow-colored stools. Probiotic- or prebiotic-treated groups did not show appreciable improvement in stool consistency; stools in these groups remained soft to semi-formed and often yellowish, indicating only partial protection. In contrast, prophylactic administration of synbiotics completely prevented the diarrhoeal phenotype; the stools of synbiotic-fed mice remained firm and brown, similar to those of uninfected controls (Figure 2B).

**Figure 2:**
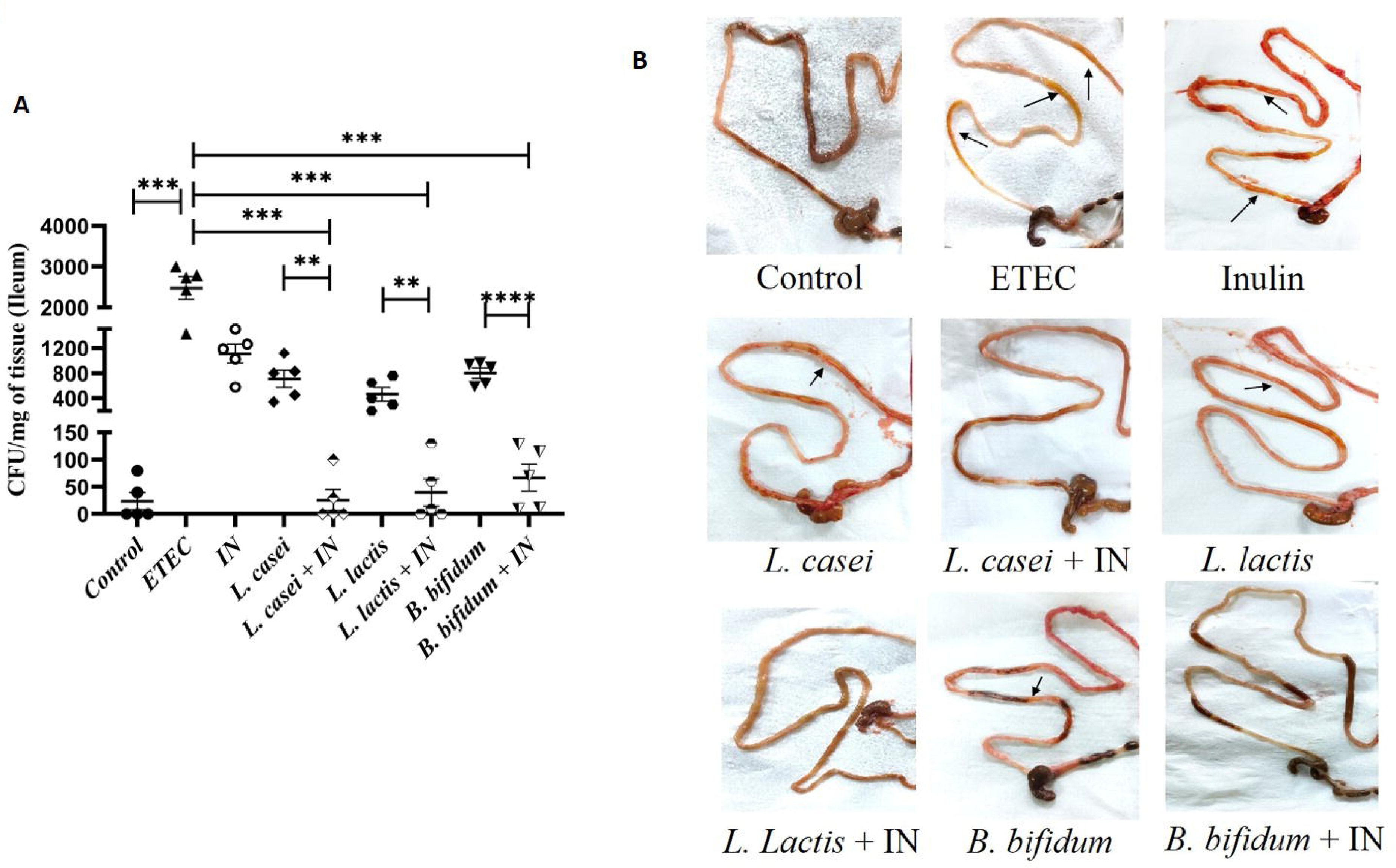
(A) 4-6-week-old mice were grouped into control, ETEC, IN, *L. casei*, *L. casei*+IN, *L. lactis*, *L. lactis* +IN, *B. bifidum,* and *B. bifidum* +IN. Apart from the control, all groups were infected with ETEC H10407 at 10^9^ CFU/ml. To assess the colonization of ETEC H10407, small ileal sections were isolated from each group and homogenized in PBS and then the lysates were spread onto the MacConkey agar plate. Infection loads in each and every group were compared with the ETEC group. And each synbiotic group was compared with the corresponding probiotic and IN group. (B) Representative images of stool consistency and intestinal fluid accumulation are shown. ETEC-infected mice exhibited watery, yellow stools with pronounced fluid accumulation (black arrows). Probiotic- and prebiotic-treated groups displayed partial protection, with stools remaining soft to semi-formed and yellowish. In contrast, prophylactic synbiotic treatment completely prevented the diarrheal phenotype, with stools remaining firm and brown, comparable to uninfected controls. The images of small intestinal contents of mice from different experimental groups. Experiments were conducted with n ≥ 5 animals per group, and the data are represented as Mean ± SEM. * represents p < 0.05, ** represents p < 0.01, *** represents p < 0.001, **** represents p <0.0001.

### Histological analysis of colonic tissue sections

Histological studies were used to examine tissue injury and inflammation in order to study the change in the cellular morphology of the ileum of the small intestine. Figure 3A presents the findings. The following are listed in that order: (a) IN, (b) ETEC, and (c) Control. (d) *L. casei*, (e) *L. lactis*, (f) *B. bifidum*, (g) *L. casei* + IN, (h) *L. lactis* + IN, and (I) *B. bifidum* + IN. While the animals in the control group displayed intact mucosa, tubular crypts, and goblet cells with no detectable scattered immune cell infiltration at the crypt and mucosal layers, the ETEC group displayed a significant loss of crypts and goblet cells.

**Figure 3:**
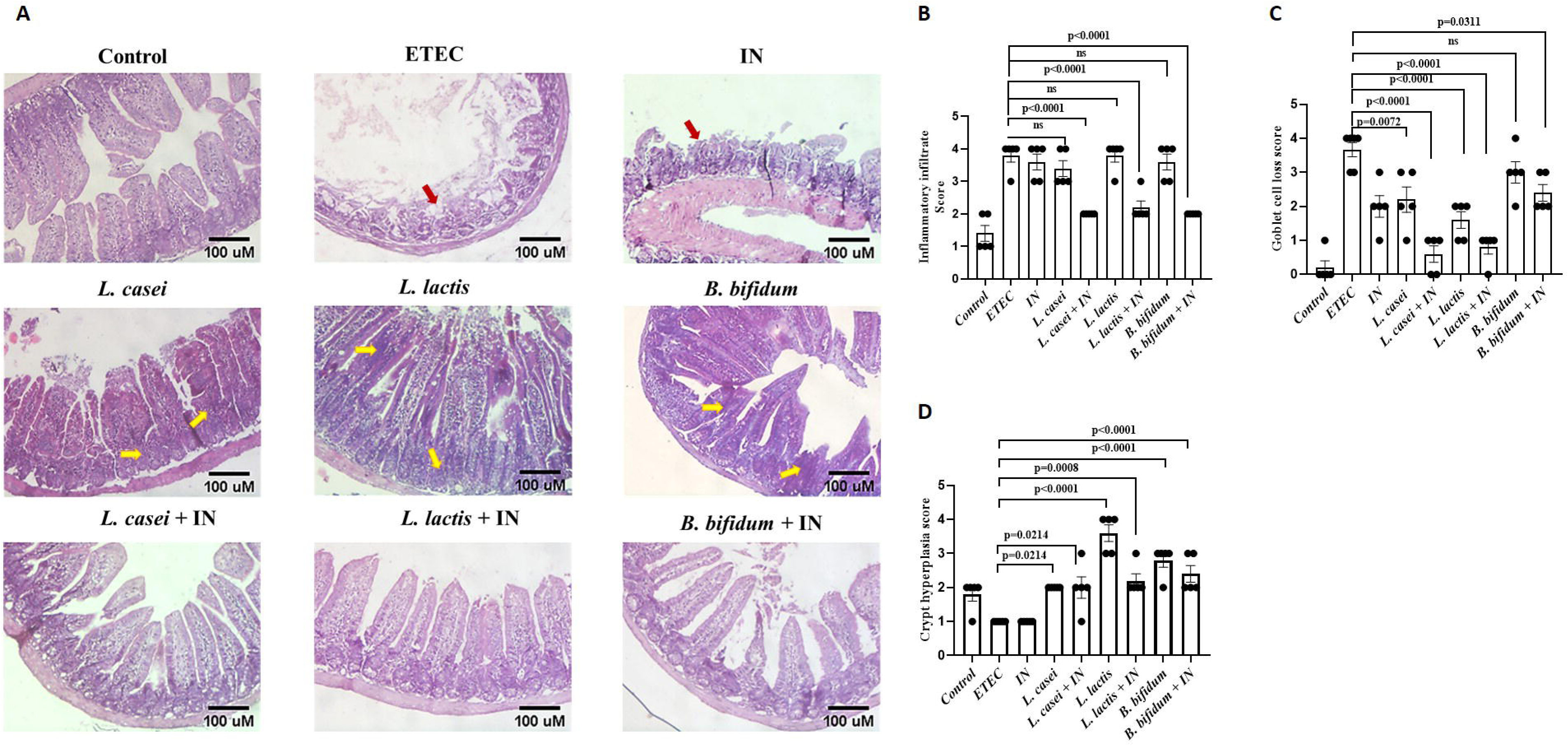
(A) Histological examination of the cross sections of the ileum of mice from different groups. Tissues were stained with Haematoxylin and Eosin and viewed under the microscope with 40X magnification to understand the tissue histology. A Histological score was generated according to the degree of changes in the (B) Inflammatory infiltrate, (C) Goblet cell loss, and (D) Crypt hyperplasia, and compared among the groups.

**Figure 4:**
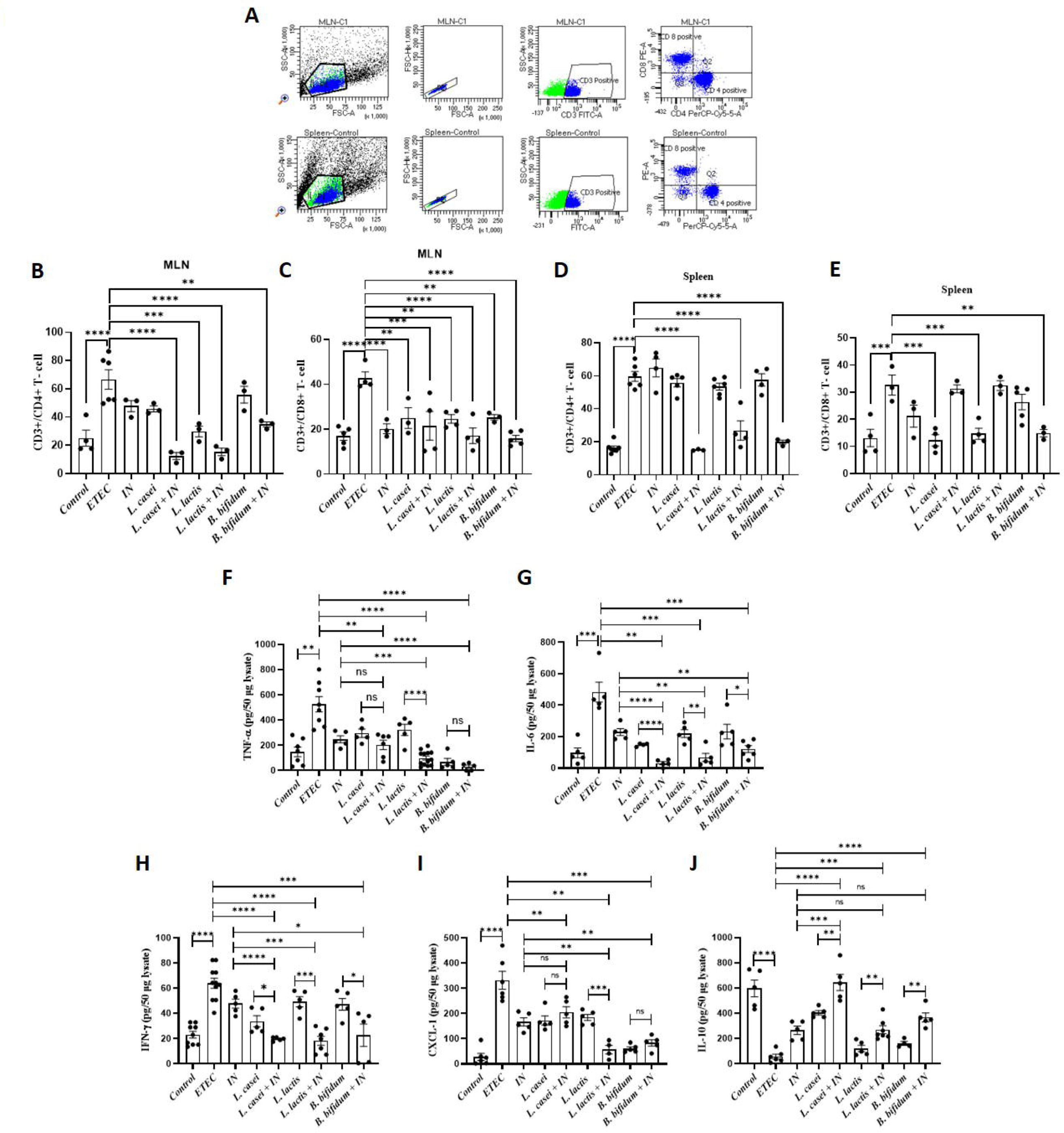
(A) Scattered plot for the gating strategy of CD3+CD4+ and CD3+CD8+ cell acquisition by flow cytometry from the cells of MLN and spleen of experimental mice groups. The singlet population of the parent population was scattered according to their expression of CD4+CD8-(Q4), CD4-CD8+(Q1), CD4+CD8+(Q2), and CD4-CD8-(Q3). (B-E) The population for CD4+ and CD8+ T cells in both MLN and spleen was determined by flow cytometry. Cells harvested from both MLN and spleen were tagged with anti-CD3, anti-CD4, and anti-CD8 antibodies. CD4+ and CD8+ cells were detected from the CD3+ parent population of T cells. (F-J) Determination of pro-inflammatory markers such as TNF-α, IL-6, IFN-γ, CXCL-1, and anti-inflammatory IL-10 expression by ELISA from ileal tissue lysates. 50µg protein lysates from each sample were subjected to ELISA. Cytokine expressions of each synbiotic group were compared with the corresponding probiotic group, prebiotic or IN group, and ETEC group. N ≥ 5 animals were taken per group, and the data are represented as Mean ± SEM. * represents p < 0.05, ** represents p < 0.01, *** represents p < 0.001, **** represents p <0.0001.

**Figure 5:**
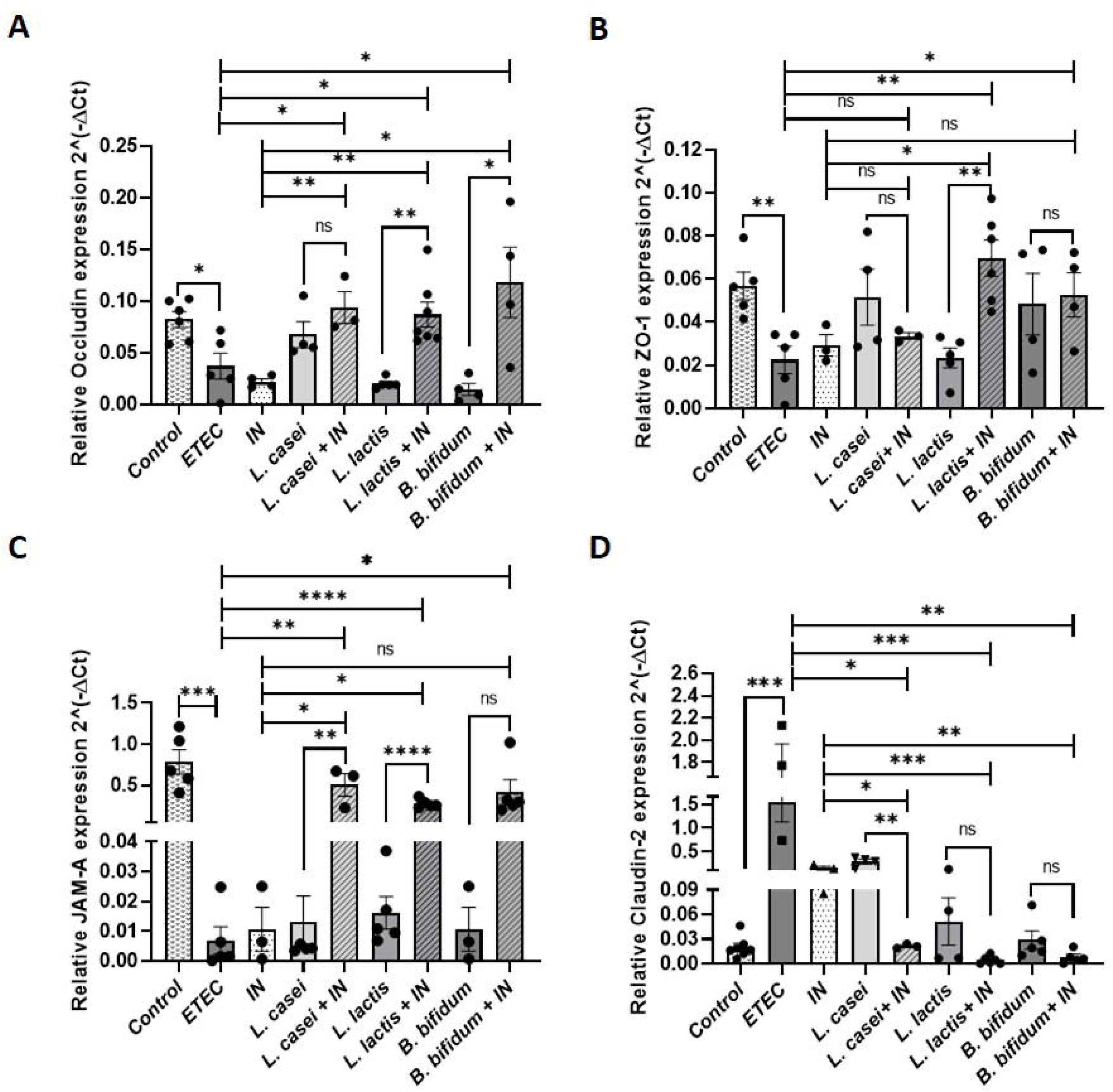
Epithelial tight junctional barrier integrity was analysed by determining the mRNA expression of Occludin, ZO-1, and JAM-A proteins by real-time PCR. mRNA expression of another tight junctional protein, Claudin-2, which is a pore-forming protein, was also detected by real-time PCR. GAPDH was set as the internal control mRNA. Data was plotted using the relative gene expression values. Gene expressions of tight junction proteins of each synbiotic group were compared with the corresponding probiotic group, prebiotic or IN group, and ETEC group. This study was conducted with N ≥ 3 animals per group, and the data are represented as Mean ± SEM * represents p < 0.05, ** represents p < 0.01, *** represents p < 0.001, **** represents p <0.0001.

**Figure 6:**
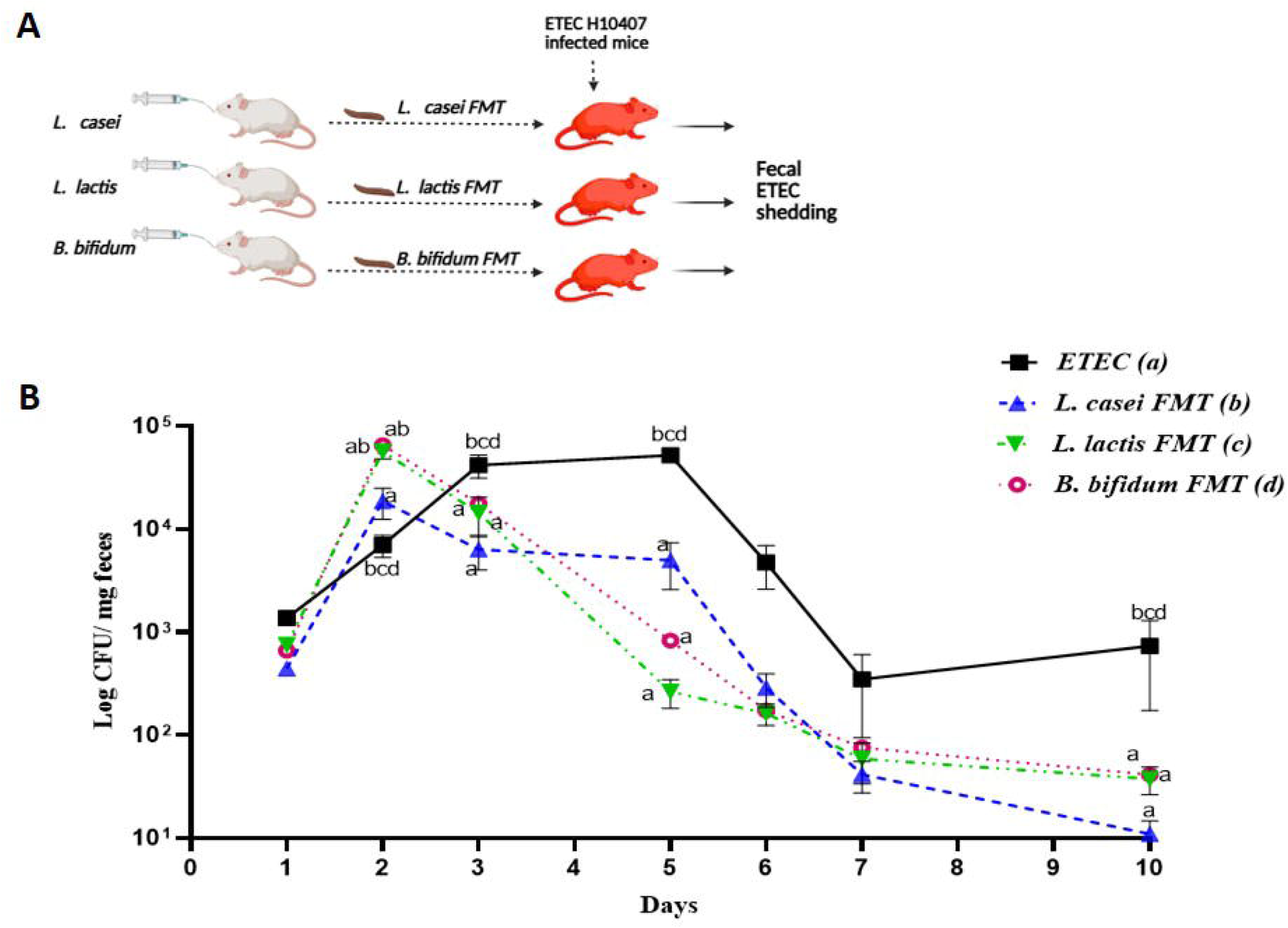
Day-wise monitoring of the fecal ETEC CFU load. Pathogens (CFU) are excreted with fecal pellets after infection. Data were collected at different time intervals. Error bars indicate sem.

The ETEC group showed a 55.55% reduction in crypt length, significantly less than that of the control group (P < 0.0001). Crypt hyperplasia was noticed in *L. lactis*, *B. bifidum*, compared to the control, which was a 2 and 1.55-fold increase, respectively. But in *L. lactis + IN,* a 22%-or 1.22-fold increase and in *B. bifidum + IN* groups, a 33.33%-or 1.33-fold increase in crypt length was observed compared to the control crypt length, which was not a significant change. The ileal crypt structure was less altered in the *L. casei,* and *L. casei + IN* group; the mucosa and submucosa layers remained nearly intact, but the cells inside this layer were flaccid, and neutrophil infiltration was noted in these layers. When compared to the ETEC group, the immune infiltration was significantly (p<0.0001) less in IN, *L. casei, L. casei + IN*, *L. lactis, L. lactis + IN, B. bifidum,* and *B. bifidum + IN* groups, with a damage score of 5.26%, 15.23%, 55.39%, 55.38%, 74.17%, 75.53% and 87% less than ETEC. This tells that the *L. casei + IN* showed a 40% improvement over the *L. casei*, *L. lactis + IN* had an 18.78% improvement over the *L. lactis group,* and *B. bifidum + IN* showed an 11.59% improvement over *B. bifidum.* Looking into the goblet cell loss in the ileal crypts, a significant (p<0.0001) 95% reduction of goblet cells was observed in the ETEC ileal sections when compared to the control. However, the prebiotic and probiotic groups IN, *L. casei, L. lactis*, and *B. bifidum,* exhibited a 55%, 61%, 48.6%, and 83.33% reduction in disease score, respectively. The disease score for goblet cell loss in the synbiotic groups *L. casei* + IN, *L. lactis* + IN, and *B. bifidum* + IN showed a further reduction of 47.22%, 26.39%, and 16.67%, respectively (Figure 3B).

### Synbiotics reduce the CD3+CD4+ Th1 cell population in ETEC-induced inflammation

In association with the cytokine profile, this study investigated the T-cell population of the spleen and mesenteric lymph nodes (MLN) in each test animal after 20 hours post-infection. In the spleen, the CD3+CD4+ T-cells were increased 3.21-fold in the ETEC group compared to the control, with a p value of <0.0001. CD3+CD4+ T-cells showed a significant population decrease in all the synbiotic groups, which was 47.48% (p= 0.0002) in *L. casei* + IN group, 54.9% (p= 0.0001) in the *L. lactis* + IN group and 66.83% (p <0.0001) in the *B. bifidum* + IN group, compared to the ETEC group. However, no significant change was observed in the IN, *L. casei*, *L. lactis,* and *B. bifidum* prebiotic and probiotic groups compared to the ETEC group. Also, *L. casei* + IN, *L. lactis* + IN and *B. bifidum* + IN showed 40.95% (p= 0.0033), 44.51% (p= 0.0026) and 63.19% (p= 0.0002) decrease in CD3+CD4+ T-cell population when compared with respective probiotic groups and a decrease of 56.11% (p<0.0001), 63.52% (p<0.0001) and 75.46% (p<0.0001) when compared with IN group. ETEC-infected mice exhibited a significantly increased number of splenic CD3+CD8+ T cells, representing a 60% increase compared to the control mice. In comparison to the ETEC infection group, the CD3+CD8+ T cell percentage among the therapeutic groups showed a decrease of 35.14% in IN, 62.4% in *L. casei*, 55% in *L. lactis,* and 19.16% in *B. bifidum* groups. However, no significant change was observed in both *L. casei*+ IN.

### Prebiotics, probiotics, and synbiotics reduce inflammatory cytokines in the ileal milieu

In this study, we also investigated the state of inflammation upon ETEC colonization in the intestinal tissues. The major pro-inflammatory cytokines, such as TNF-α, IL-6, IFN-γ, and anti-inflammatory IL-10 secretion, were assessed by ELISA from the colonic tissue lysates of each sample. In the ETEC group, the tissue TNF-α (pg/ml) 526.19 ± 173.31, IL-6 (pg/ml) 482.97 ± 142.005, and IFN-γ (pg/ml) 64.04 ± 13.18 were increased compared to the control group, which was 99.816 ± 71.037, 97.064 ± 70.63, and 23.12 ± 8.056, respectively. This data indicates a significant upregulation of TNF-α, IL-6, and IFN-γ in the ETEC group with a change in the fold of 5.27 (p<0.0001), 4.976 (p<0.0001), and 2.77 (p<0.0001), respectively, compared to the control cytokine levels. TNF-α expression in IN, *L. casei*, *L. lactis,* and *B. bifidum* was 248 ± 61.07, 296.53 ± 74.27, 322.44 ± 99.49, and 68.67 ± 38.16, which indicates a reduction of 52.87% (p= 0.0001), 43.64% (p= 0.0027), 38% (p= 0.0116), and 86% (p <0.0001) from the ETEC group. Surprisingly, in the *L. casei* + IN, *L. Lactis*+ IN, and *B. bifidum*+ IN groups the TNF-α expression was 202.83 ± 92.45, 141.54 ± 34.73, and 49.6 ± 12.04, having further reduction of 17.8% (p= 0.7808), 34.38% (p= 0.0013), and 3.62% (p= 0.9992) from respective probiotic groups. Compared to the ETEC group, IL-6 expression was significantly decreased to 52.52% (p= 0.0001) in the IN, 69.24% (p <0.0001) in the *L. casei*, 54.56% (p <0.0001) in the *L. lactis* and 51.91% (p= 0.0001) in the *B. bifidum* group. Compared to the respective probiotic groups, we found that IL-6 production was further decreased with a percentage of 24.15% (p<0.0001), 31.43% (p= 0.0029), and 22.95% (p= 0.0471) in *L. casei* + IN, *L. Lactis*+ IN, and *B. bifidum*+ IN synbiotic groups, respectively. Similarly, with a decrease of 25.24% (p= 0.1663), 47.18% (p= 0.0002), 22.9% (p= 0.2709), and 26.17% (p= 0.1346) respectively, IN, *L. casei*, *L. lactis,* and *B. bifidum* groups showed a significant decrease in IFN-γ production compared to the ETEC group. Having similarities with the previous data, we found even more decrease in IFN-γ in the *L. casei* + IN, *L. Lactis* + IN, and *B. bifidum* + IN synbiotic groups, which were 21% (p= 0.0125), 48% (p= 0.0005), and 38% (p= 0.0236) compared to their respective probiotic groups. On the other hand, IL-10, as the anti-inflammatory cytokine, which was 597.683 ± 148.219 in the control group, was significantly (p<0.0001) decreased to 66.35 ± 43.67 in the ETEC group, showing a reduction by 88.89% or a 9-fold decrease compared to the control. From this experiment, we got a significant increase in IL-10 secretion in IN, *L. casei*, *L. lactis* and *B. bifidum* groups, which is 74.95% (p= 0.0092), 83.58% (p <0.0001), 45.39% (p = 0.9455), and 58.9% (p= 0.6435), respectively. Further increase in IL-10 production was seen in the *L. casei* + IN, *L. Lactis* + IN and *B. bifidum* + IN synbiotic groups, which is 6.12% (p= 0.0067), 30.035% (p= 0.0060) and 23.05% (p= 0.0020), respectively, compared to their respective probiotic groups. Besides all the inflammatory cytokines we investigated the expression of CXCL-1 production in the ileal tissues which is a major neutrophil chemoattractant. In the ETEC group the CXCL-1 level was 332.183 ± 86.748 which is an 11.76-fold increase from the control 23.1 ± 8.53. The prophylactic IN, *L. casei*, *L. lactis* and *B. bifidum* groups showed 49.75% (p<0.0001), 48.02% (p <0.0001), 44.41% (p= 0.0001) and 81.55% (p <0.0001) decrease in CXCL-1 level compared to the ETEC group. Among all three synbiotic groups, only *L. lactis* + IN group inhibited 38.45% (p= 0.0047) more reduction in CXCL-1 expression when compared with its corresponding probiotic group.

### Restoration of tight junction proteins by prebiotic, probiotic, and synbiotic

This study investigated major tight junction proteins associated with epithelial cells, including transmembrane Occludin, claudins, and cytoplasmic protein ZO-1. A 2.34-fold decrease in the Occludin mRNA expression was observed in the ETEC H10407-infected ileal tissues compared to the control ileum. Among IN, *L. casei*, *L. lactis,* and *B. bifidum* prebiotic and probiotic groups, only the *L. casei* group showed an upregulation of Occludin with an increase of 1.8-fold. Occludin expression was significantly upregulated in all three synbiotic groups with an increase of 2.53, 2.35, and 3.18 folds or 60.43%, 57.4% and 68.63% respectively when compared to the ETEC group (Figure-5A).

ZO-1 was seen to be decreased 2.53-fold or 60.47% in the ETEC group compared to the control group. *L. casei* probiotic group, as well as *L. casei* + IN synbiotic group, showed no significant improvement in the ZO-1 upregulation. Not the *L. lactis* probiotic group but the *L. lactis* + IN synbiotic group exhibited a 3.1-fold or 67.8% increase. Interestingly, the *B. bifidum* probiotic and *B. bifidum* + IN synbiotic group both showed significant upregulation of ZO-1 with an increase of 53.59% and 57.45% respectively (Figure-5B). A 99% reduction of JAM-A expression was observed in ETEC-infected mice. We found no significant upregulation of JAM-A in all three probiotic groups, along with the IN group. On the contrary, all three synbiotic groups showed a significant increase of 98.63% for *L. casei* + IN, 97.54% for *L. lactis* + IN, and 98.35% for *B. bifidum* + IN, respectively (Figure-5C). Claudin-2, the pore-forming claudin family TJ protein, was significantly increased by 76.43% or 128-fold in ETEC-infected mice; however, a significant improvement was observed in the IN, *L. casei*, *L. lactis,* and *B. bifidum* groups, where the Claudin-2 expression was 9.4%, 18.35%, 3.32% and 1.23% respectively, relative to the ETEC group. Notably, all synbiotic groups showed a further and more pronounced suppression of Claudin-2 expression: 1.2% in *L. casei* + IN, 0.39% in *L. lactis* + IN, and 0.45% in *B. bifidum* + IN, which were statistically comparable to the control group, suggesting that synbiotic treatment was most effective in restoring Claudin-2 to baseline levels and thereby preserving epithelial barrier integrity (Figure-5D).

### Probiotic-induced gut microbiome alterations help ETEC clearance

In search for the beneficial effect of Probiotic consumption-mediated Gut microbial alteration, and its protection against ETEC colonization, the fecal pellets of probiotic-fed mice were collected and orally given to ETEC-infected mice. The kinetics of pathogen clearance among ETEC vs probiotic FMT groups were similar on day one. On day two, pathogen shedding was visibly increased in the probiotic-FMT-grouped mice compared to the ETEC group. A subsequent sharp decline in ETEC shedding was observed among the probiotic FMT groups from day 3 and onwards (Figure-6), while in the ETEC group, a significant (p< 0.0001) decline in ETEC load within fecal matter was observed on day 6 and onward.

## Discussion

### ETEC Infection: The Need for Alternative Prophylactics

Enterotoxigenic Escherichia coli (ETEC) remains one of the most common causes of traveler’s diarrhoea, particularly in developing countries. Despite the efficacy of antibiotics like fluoroquinolones in healthy individuals, prophylactic use is not advised due to the risk of developing antibiotic-resistant strains (55) (56) (57). Again, the risk of generating an antibody response against ETEC toxin neutralizers keeps the problem unsolved (58). Additionally, vaccine development has faced challenges, including concerns over inadequate immunogenicity and potential autoimmune responses due to antibody cross-reactivity with host tissues (59), (60). As such, safe, accessible, and effective non-antibiotic alternatives are urgently needed.

In this context, our study explores probiotic and synbiotic-based prophylactic interventions using *Lactobacillus casei*, *Lactococcus lactis*, and *Bifidobacterium bifidum*—alone and in combination with inulin—against ETEC H10407-induced intestinal inflammation in a murine model.

### Probiotics and Synbiotics Inhibit ETEC Colonization and Adhesion

From the secret goat yogurt reportedly used by a Jewish Ottoman doctor in the 16th century to treat France’s King Francis I for a severe gastrointestinal illness, to Pasteur’s 1857 discovery of lactic acid-producing bacteria, the therapeutic use of probiotics has evolved significantly (61). Today, probiotics are recognized as cutting-edge tools for both prevention and treatment of various health conditions. The genera *Lactobacillus* and *Bifidobacterium* comprise the most commonly utilized probiotic species (62), (63). Specifically, *L. casei*, *L. lactis*, and *B. bifidum* have demonstrated probiotic efficacy against infectious diarrhoea caused by the LT⁺ ST⁺ ETEC H10407 strain in murine models, with reduced colonization observed in the ileal segment of the small intestine following prophylactic administration. Expanding on these findings, our data demonstrate that all three probiotic strains significantly reduced ETEC growth and adhesion in vitro, an effect further enhanced when combined with inulin to form synbiotics. Adhesion inhibition and competitive exclusion assays using INT 407 cells revealed a highly significant reduction in ETEC adherence (p < 0.0001) after pre-treatment with the probiotic strains, highlighting their potential in preventing initial colonization. In vivo, mice receiving synbiotic treatment exhibited a greater reduction in intestinal ETEC load than those treated with probiotics alone, confirming a synergistic effect. The addition of inulin to each probiotic strain significantly decreased bacterial colonization in the small intestine, improved bacterial clearance kinetics, and reduced the severity of diarrhoeal symptoms.

### Histological Protection and Goblet Cell Preservation

Histological analysis of ileal tissues revealed that ETEC infection resulted in crypt loss, goblet cell depletion, and significant immune cell infiltration. Synbiotic-treated groups showed preserved mucosal architecture and reduced inflammatory cell infiltration compared to the ETEC group. Specifically, the *L. casei* + IN group showed a 40% improvement in damage score over *L. casei* alone, while the *L. lactis* + IN and *B. bifidum* + IN groups exhibited 18.78% and 11.59% improvements, respectively. Goblet cell depletion, a key marker of epithelial damage, was significantly ameliorated in synbiotic-treated groups, further supporting their role in maintaining mucosal integrity.

### Synbiotics Attenuate Inflammatory T-Cell Responses

The observed reduction in CD3⁺CD4⁺ Th1 and CD3⁺CD8⁺ T-cell populations in synbiotic-treated groups suggests that synbiotics provide a more effective immunoregulatory response to ETEC infection compared to probiotics or inulin alone. This enhanced effect may be due to improved colonization and stability of *Lactobacillus species* in the gut when supported by inulin. *Lactobacilli* not only metabolize bile salts and synthesize essential vitamins B and K, but also contribute to the maturation of both innate and adaptive immune responses. Furthermore, their role in suppressing pro-inflammatory mediators (64) underscores their potential in mitigating gut inflammation and maintaining immune homeostasis during enteric infections.

### Cytokine Modulation and Reduced Chemokine Expression

*Enterotoxigenic Escherichia coli* (ETEC) infection induces a robust inflammatory response characterized by elevated pro-inflammatory cytokines such as TNF-α, IL-6, and IFN-γ, alongside increased levels of the chemokine CXCL-1. This heightened inflammatory milieu is coupled with a significant downregulation of the anti-inflammatory cytokine IL-10, contributing to tissue inflammation and immune dysregulation. Our data demonstrate that synbiotic treatment—particularly the combination of *Lactococcus lactis* and inulin—effectively reverses these trends more than probiotic or prebiotic treatments alone. This synbiotic formulation notably reduced CXCL-1 expression, correlating with decreased neutrophil infiltration, and enhanced IL-10 levels, highlighting its superior immunomodulatory capacity.

These findings align with observations in human ETEC infections, where severe colonization correlates with increased myeloperoxidase (MPO) levels in stool, indicative of systemic inflammation. Elevated serum IL-17A and IFN-γ in infected volunteers further support the role of these cytokines as markers of heightened inflammatory activity (65). The increased neutrophil presence observed in the ileal villi of infected mice mirrors the elevated MPO expression found in human cases. Moreover, the outer membrane vesicles (OMVs) of ETEC have been shown to potently induce pro-inflammatory responses; notably, wild-type ETEC H10407 stimulates IL-8 secretion in human colon epithelial cells (HT-29) (66), paralleling the upregulation of CXCL-1 observed in our murine model. Given the functional homology between IL-8 in humans and CXCL-1 in mice as neutrophil chemoattractants (67), these results suggest conserved mechanisms driving neutrophil recruitment and inflammation across species. Together, these data support the therapeutic potential of synbiotic interventions in mitigating ETEC-induced intestinal inflammation by modulating both pro- and anti-inflammatory cytokine profiles and reducing neutrophil-mediated tissue damage.

### Restoration of Tight Junction Integrity

Previous studies have demonstrated that probiotics can alleviate antibiotic-associated diarrhoea (AAD) and restore gut microbial balance by upregulating tight junction (TJ) proteins such as occludin, ZO-1, and claudin-1. This effect was accompanied by an increased abundance of beneficial bacterial genera including *Lactobacillus*, *Bacteroides*, and *Muribaculaceae* (68). Similar protective effects were reported in an intestinal multidrug-resistant *Acinetobacter baumannii* (MDRAb) infection model, where the synbiotic combination of the prebiotic galactooligosaccharides (GOS) and the probiotic *Bifidobacterium breve* strain Yakult (BbY) improved gut integrity and immune responses (69).

In addition, *Lactobacillus casei* ATCC 393 was shown to protect IPEC-J2 cells against ETEC K88-induced loss of transepithelial electrical resistance by restoring TJ proteins ZO-1 and occludin. In porcine mast cell cultures, ETEC K88 reduced anti-inflammatory cytokines and increased pro-inflammatory mediators including IL-1β, β-hexosaminidase, tryptase, TNF-α, and IFN-γ. Pretreatment with *L. casei* ATCC 393 mitigated these inflammatory responses and preserved protein expression (70).

Our findings provide further evidence of the protective role of *Lactobacillus casei* ATCC 393 against ETEC H10407-induced intestinal barrier dysfunction and inflammation. Specifically, we observed that synbiotic treatment with *L. casei* ATCC 393 effectively restored the expression of critical tight junction (TJ) proteins, including occludin, ZO-1, and JAM-A, which are essential for maintaining epithelial integrity and preventing paracellular permeability. The restoration of these TJ proteins likely contributed to the significant reduction in claudin-2 overexpression, a protein commonly associated with increased epithelial permeability and barrier disruption during enteric infections. Moreover, oral vaccination with *Lactococcus lactis* has demonstrated protective efficacy against LT-producing ETEC in a rabbit model by inhibiting fluid loss (71). To our knowledge, this is the first evidence supporting *Lactococcus lactis* ATCC 49032 as a promising probiotic against ETEC-induced diarrhoea.

### Gut Microbiota Transfer Confirms Probiotic-Mediated Protection

This study demonstrates that fecal microbiota transplantation (FMT) from probiotic-fed mice into ETEC-infected recipients accelerated pathogen clearance. While shedding patterns were similar on day 1, probiotic-FMT mice exhibited a transient rise in ETEC on day 2, followed by a sharp decline from day 3 onward. In contrast, ETEC-only mice showed a delayed reduction beginning on day 6. The transient increase in shedding on day 2 may represent an ecological adjustment as transplanted microbes competed with resident communities and the pathogen. Such biphasic responses have been reported in FMT studies, where initial perturbations precede stable pathogen suppression (72). The accelerated clearance observed from day 3 highlights the protective impact of probiotic-conditioned microbiota. Probiotics can influence pathogen survival through multiple mechanisms, including production of antimicrobial metabolites, competition for epithelial binding sites, reinforcement of barrier integrity, and immune modulation (73), (74). For example, *Lactobacillus reuteri* HCM2 has been shown to inhibit ETEC adhesion and preserve gut morphology (73).

In the ETEC-only group, clearance likely reflected the natural progression of host immunity. However, the earlier decline in the probiotic-FMT group suggests that microbiome alterations provided a competitive advantage against pathogen persistence. These results indicate that transferring a microbiota shaped by probiotic consumption can significantly reduce the duration of ETEC infection. From a translational standpoint, probiotic-conditioned FMT may provide a more robust and lasting therapeutic effect compared to probiotics alone, particularly in high-burden settings where rapid clearance of enteric pathogens is critical. Future work should focus on identifying the specific microbial taxa and metabolites responsible and clarifying host–microbe interactions underlying this protection. In conclusion, probiotic-induced microbiota alterations, when transferred via FMT, enhance ETEC clearance and represent a promising microbiome-based intervention against enteric infections.

Because travelers’ diarrhoea has a very high strike rate and makes it easy to identify those who are at risk before exposure, it is a perfect candidate for the prophylactic deployment of toxin-binding probiotics (59). As travel is generally a pre-planned event, a prophylactic administration of probiotics or synbiotics can be taken by travelers, which not only offers protection against ETEC but can prevent many other pathogenic bacteria from colonizing in the gut and can help in coping with a foreign diet.

## Conclusion

This study demonstrates that both probiotics and synbiotics offer significant prophylactic benefits against *Enterotoxigenic Escherichia coli* (ETEC) H10407-induced intestinal infection in a murine model. Probiotic strains *Lactobacillus casei*, *Lactococcus lactis*, and *Bifidobacterium bifidum* effectively reduced ETEC growth, adhesion, and colonization, and their combination with the prebiotic inulin as synbiotics further enhanced these protective effects. Synbiotic treatment resulted in a pronounced reduction in gut inflammation, preservation of intestinal architecture, restoration of tight junction proteins, and modulation of key immune responses, including reduced Th1 cell populations and pro-inflammatory cytokines. Additionally, fecal microbiota transfer from probiotic-treated mice confirmed the role of gut microbial modulation in pathogen clearance.

Taken together, these findings establish synbiotic formulations—particularly those involving *L. lactis* and *B. bifidum*—as a promising, non-antibiotic prophylactic strategy to combat ETEC infection. The synergy between probiotics and prebiotics enhances host resilience to enteric pathogens and supports mucosal immune homeostasis. Given their safety, cost-effectiveness, and ease of administration, synbiotics hold strong potential as a preventive intervention for travelers’ diarrhoea and other enteric diseases.

## Acknowledgement

The Director of ICMR-NIRBI, Kolkata, assisted the authors in conducting the study. The authors express their gratitude to Narayan Chandra Mondal for genuinely preserving the mice’s health. The central instrument facilities provided by Animesh Gope, Ananda Pal, and Biswajit Sharma are acknowledged by the authors. SS was the beneficiary of a CSIR fellowship. We acknowledge the use of ChatGPT (OpenAI) and QuillBot for language editing and paraphrasing support in preparing the manuscript. The authors confirm that all scientific content, interpretation of data, and final conclusions are solely their own.

## Author Contribution

Sohini Sikdar: Conceptualization, Methodology, Formal analysis, Investigation, Writing Original Draft, Visualization. Paulami Dutta: Methodology, Investigation, Formal analysis. Debmalya Mitra: Methodology, Resources. Bidisha Pal: Investigation. Hemanta Koley: Conceptualization, Validation, Resources, supervision. Shanta Dutta: Supervision, funding acquisition.

## Abbreviation

ETEC: Enterotoxigenic Escherichia coli
L. casei: Lactobacillus casei
L. lactis: Lactococcus lactis
B. bifidum: Bifidobacterium bifidum
CFU: Colony Forming Unit
MOI: Multiplicity of infection
IN: Inulin
LT: Heat-Labile Toxin
ST: Heat Stable Toxin
MLN: Mesenteric Lymph Node
IL: Interleukin
TNF: Tumor Necrosis Factor
IFN: Interferon
TJ: Tight Junction
ZO-1: Zonula Occludens-1
JAM-A: Junctional Adhesion Molecule-A
FMT: Fecal Microbiota Transplantation

## References

1. Qadri F, Saha A, Ahmed T, Al Tarique A, Begum YA, Svennerholm AM. Disease burden due to enterotoxigenic Escherichia coli in the first 2 years of life in an urban community in Bangladesh. Infect Immun. 2007;75(8):3961–8.

2. Ashkenazi S, Schwartz E, O’Ryan M. Travelers’ Diarrhea in Children: What Have We Learnt? Pediatr Infect Dis J. 2016;35(6):698–700.

3. Steffen R, Hill DR, DuPont HL. Traveler’s diarrhea: a clinical review. JAMA. 2015;313(1):71–80.

4. Stoney RJ, Han PV, Barnett ED, Wilson ME, Jentes ES, Benoit CM, et al. Travelers’ Diarrhea and Other Gastrointestinal Symptoms Among Boston-Area International Travelers. Am J Trop Med Hyg. 2017;96(6):1388–93.

5. DuPont HL. Systematic review: the epidemiology and clinical features of travellers’ diarrhoea. Aliment Pharmacol Ther. 2009;30(3):187–96.

6. Leung AKC, Leung AAM, Wong AHC, Hon KL. Travelers’ Diarrhea: A Clinical Review. Recent Pat Inflamm Allergy Drug Discov. 2019;13(1):38–48.

7. Madhavan TP, Sakellaris H. Colonization factors of enterotoxigenic Escherichia coli. Adv Appl Microbiol. 2015;90:155–97.

8. Zhang Y, Tan P, Zhao Y, Ma X. Enterotoxigenic Escherichia coli: intestinal pathogenesis mechanisms and colonization resistance by gut microbiota. Gut Microbes. 2022;14(1):2055943.

9. Svennerholm AM, Lundgren A. Developments in oral enterotoxigenic Escherichia coli vaccines. Curr Opin Immunol. 2023;84:102372.

10. Khalil I, Walker R, Porter CK, Muhib F, Chilengi R, Cravioto A, et al. Enterotoxigenic Escherichia coli (ETEC) vaccines: Priority activities to enable product development, licensure, and global access. Vaccine. 2021;39(31):4266–77.

11. Anderson JDt, Bagamian KH, Muhib F, Baral R, Laytner LA, Amaya M, et al. Potential impact and cost-effectiveness of future ETEC and Shigella vaccines in 79 low- and lower middle-income countries. Vaccine X. 2019;2:100024.

12. Bourgeois AL, Wierzba TF, Walker RI. Status of vaccine research and development for enterotoxigenic Escherichia coli. Vaccine. 2016;34(26):2880–6.

13. Tribble DR. Resistant pathogens as causes of traveller’s diarrhea globally and impact(s) on treatment failure and recommendations. J Travel Med. 2017;24(suppl_1):S6–S12.

14. Li S, Upadhyay I, Seo H, Vakamalla SSR, Madhwal A, Sack DA, et al. Immunogenicity and preclinical efficacy characterization of ShecVax, a combined vaccine against Shigella and enterotoxigenic Escherichia coli. Infection and Immunity.0(0):e00004-25.

15. Gaastra W, Svennerholm AM. Colonization factors of human enterotoxigenic Escherichia coli (ETEC). Trends Microbiol. 1996;4(11):444–52.

16. Wolf MK. Occurrence, distribution, and associations of O and H serogroups, colonization factor antigens, and toxins of enterotoxigenic Escherichia coli. Clin Microbiol Rev. 1997;10(4):569–84.

17. Qadri F, Svennerholm AM, Faruque AS, Sack RB. Enterotoxigenic Escherichia coli in developing countries: epidemiology, microbiology, clinical features, treatment, and prevention. Clin Microbiol Rev. 2005;18(3):465–83.

18. Kantele A, Riekkinen M, Jokiranta TS, Pakkanen SH, Pietila JP, Patjas A, et al. Safety and immunogenicity of ETVAX(R), an oral inactivated vaccine against enterotoxigenic Escherichia coli diarrhoea: a double-blinded, randomized, placebo-controlled trial amongst Finnish travellers to Benin, West Africa. J Travel Med. 2023;30(7).

19. Hossain MJ, Svennerholm AM, Carlin N, D’Alessandro U, Wierzba TF. A Perspective on the Strategy for Advancing ETVAX((R)), An Anti-ETEC Diarrheal Disease Vaccine, into a Field Efficacy Trial in Gambian Children: Rationale, Challenges, Lessons Learned, and Future Directions. Microorganisms. 2023;12(1).

20. Damaskos D, Kolios G. Probiotics and prebiotics in inflammatory bowel disease: microflora ’on the scope’. Br J Clin Pharmacol. 2008;65(4):453–67.

21. Li C, Peng K, Xiao S, Long Y, Yu Q. The role of Lactobacillus in inflammatory bowel disease: from actualities to prospects. Cell Death Discov. 2023;9(1):361.

22. Qin S, Huang Z, Wang Y, Pei L, Shen Y. Probiotic potential of Lactobacillus isolated from horses and its therapeutic effect on DSS-induced colitis in mice. Microb Pathog. 2022;165:105216.

23. Slizewska K, Markowiak-Kopec P, Slizewska W. The Role of Probiotics in Cancer Prevention. Cancers (Basel). 2020;13(1).

24. Reis SAD, da Conceicao LL, Peluzio M. Intestinal microbiota and colorectal cancer: changes in the intestinal microenvironment and their relation to the disease. J Med Microbiol. 2019;68(10):1391–407.

25. Liao W, Chen C, Wen T, Zhao Q. Probiotics for the Prevention of Antibiotic-associated Diarrhea in Adults: A Meta-Analysis of Randomized Placebo-Controlled Trials. J Clin Gastroenterol. 2021;55(6):469–80.

26. Guo Q, Goldenberg JZ, Humphrey C, El Dib R, Johnston BC. Probiotics for the prevention of pediatric antibiotic-associated diarrhea. Cochrane Database Syst Rev. 2019;4(4):CD004827.

27. Abdelhalim MM, Saafan GS, El-Sayed HS, Ghaith DM. In vitro antibacterial effect of probiotics against Carbapenamase-producing multidrug-resistant Klebsiella pneumoniae clinical isolates, Cairo, Egypt. J Egypt Public Health Assoc. 2022;97(1):19.

28. Iqbal Z, Ahmed S, Tabassum N, Bhattacharya R, Bose D. Role of probiotics in prevention and treatment of enteric infections: a comprehensive review. 3 Biotech. 2021;11(5):242.

29. Brussow H. Probiotics and prebiotics in clinical tests: an update. F1000Res. 2019;8.

30. Ding YH, Qian LY, Pang J, Lin JY, Xu Q, Wang LH, et al. The regulation of immune cells by Lactobacilli: a potential therapeutic target for anti-atherosclerosis therapy. Oncotarget. 2017;8(35):59915–28.

31. Corr SC, Gahan CG, Hill C. Impact of selected Lactobacillus and Bifidobacterium species on Listeria monocytogenes infection and the mucosal immune response. FEMS Immunol Med Microbiol. 2007;50(3):380–8.

32. Wang C, Li W, Wang H, Ma Y, Zhao X, Zhang X, et al. Saccharomyces boulardii alleviates ulcerative colitis carcinogenesis in mice by reducing TNF-alpha and IL-6 levels and functions and by rebalancing intestinal microbiota. BMC Microbiol. 2019;19(1):246.

33. Kobyliak N, Abenavoli L, Mykhalchyshyn G, Kononenko L, Boccuto L, Kyriienko D, et al. A Multi-strain Probiotic Reduces the Fatty Liver Index, Cytokines and Aminotransferase levels in NAFLD Patients: Evidence from a Randomized Clinical Trial. J Gastrointestin Liver Dis. 2018;27(1):41–9.

34. Ianiro G, Rizzatti G, Plomer M, Lopetuso L, Scaldaferri F, Franceschi F, et al. Bacillus clausii for the Treatment of Acute Diarrhea in Children: A Systematic Review and Meta-Analysis of Randomized Controlled Trials. Nutrients. 2018;10(8).

35. Guarino MPL, Altomare A, Emerenziani S, Di Rosa C, Ribolsi M, Balestrieri P, et al. Mechanisms of Action of Prebiotics and Their Effects on Gastro-Intestinal Disorders in Adults. Nutrients. 2020;12(4).

36. Camilleri M. Human Intestinal Barrier: Effects of Stressors, Diet, Prebiotics, and Probiotics. Clin Transl Gastroenterol. 2021;12(1):e00308.

37. Wang J, Ji H, Wang S, Liu H, Zhang W, Zhang D, et al. Probiotic Lactobacillus plantarum Promotes Intestinal Barrier Function by Strengthening the Epithelium and Modulating Gut Microbiota. Front Microbiol. 2018;9:1953.

38. You S, Ma Y, Yan B, Pei W, Wu Q, Ding C, et al. The promotion mechanism of prebiotics for probiotics: A review. Front Nutr. 2022;9:1000517.

39. Stinson LF, Payne MS, Keelan JA. Planting the seed: Origins, composition, and postnatal health significance of the fetal gastrointestinal microbiota. Crit Rev Microbiol. 2017;43(3):352–69.

40. Nyman M. Fermentation and bulking capacity of indigestible carbohydrates: the case of inulin and oligofructose. Br J Nutr. 2002;87 Suppl 2:S163–8.

41. van der Beek CM, Canfora EE, Kip AM, Gorissen SHM, Olde Damink SWM, van Eijk HM, et al. The prebiotic inulin improves substrate metabolism and promotes short-chain fatty acid production in overweight to obese men. Metabolism. 2018;87:25–35.

42. Roberfroid MB. Concepts in functional foods: the case of inulin and oligofructose. J Nutr. 1999;129(7 Suppl):1398S-401S.

43. Yang B, Lu P, Li MX, Cai XL, Xiong WY, Hou HJ, et al. A meta-analysis of the effects of probiotics and synbiotics in children with acute diarrhea. Medicine (Baltimore). 2019;98(37):e16618.

44. Schropp N, Stanislas V, Michels KB, Thriene K. How Do Prebiotics Affect Human Intestinal Bacteria?-Assessment of Bacterial Growth with Inulin and XOS In Vitro. Int J Mol Sci. 2023;24(16).

45. Wang X, Gibson GR. Effects of the in vitro fermentation of oligofructose and inulin by bacteria growing in the human large intestine. J Appl Bacteriol. 1993;75(4):373–80.

46. Buntin N, Hongpattarakere T, Ritari J, Douillard FP, Paulin L, Boeren S, et al. An Inducible Operon Is Involved in Inulin Utilization in Lactobacillus plantarum Strains, as Revealed by Comparative Proteogenomics and Metabolic Profiling. Appl Environ Microbiol. 2017;83(2).

47. Minogue TD, Daligault HA, Davenport KW, Bishop-Lilly KA, Broomall SM, Bruce DC, et al. Complete Genome Assembly of Escherichia coli ATCC 25922, a Serotype O6 Reference Strain. Genome Announc. 2014;2(5).

48. Crittenden R, Karppinen S, Ojanen S, Tenkanen M, Fagerström R, Mättö J, et al. In vitro fermentation of cereal dietary fibre carbohydrates by probiotic and intestinal bacteria. Journal of the Science of Food and Agriculture. 2002;82(8):781–9.

49. Carlson JL, Erickson JM, Hess JM, Gould TJ, Slavin JL. Prebiotic Dietary Fiber and Gut Health: Comparing the in Vitro Fermentations of Beta-Glucan, Inulin and Xylooligosaccharide. Nutrients. 2017;9(12).

50. Collado MC, Meriluoto J, Salminen S. Adhesion and aggregation properties of probiotic and pathogen strains. European Food Research and Technology. 2008:1065–73.

51. Carvalho FM, Mergulhao FJM, Gomes LC. Using Lactobacilli to Fight Escherichia coli and Staphylococcus aureus Biofilms on Urinary Tract Devices. Antibiotics (Basel). 2021;10(12).

52. Qiao H, Zhao T, Yin J, Zhang Y, Ran H, Chen S, et al. Structural Characteristics of Inulin and Microcrystalline Cellulose and Their Effect on Ameliorating Colitis and Altering Colonic Microbiota in Dextran Sodium Sulfate-Induced Colitic Mice. ACS Omega. 2022;7(13):10921–32.

53. Debmalya Mitra SS, Mainak Chakraborty, Oishika Das, Amalesh Samanta, Shanta Dutta. Gum Odina prebiotic prevents experimental colitis in C57BL/6 mice model and its role in shaping gut microbial diversity. 2023;Volume 53.

54. Erben U, Loddenkemper C, Doerfel K, Spieckermann S, Haller D, Heimesaat MM, et al. A guide to histomorphological evaluation of intestinal inflammation in mouse models. Int J Clin Exp Pathol. 2014;7(8):4557–76.

55. Riddle MS, Connor BA, Beeching NJ, DuPont HL, Hamer DH, Kozarsky P, et al. Guidelines for the prevention and treatment of travelers’ diarrhea: a graded expert panel report. J Travel Med. 2017;24(suppl_1):S57–S74.

56. Diptyanusa A, Ngamprasertchai T, Piyaphanee W. A review of antibiotic prophylaxis for traveler’s diarrhea: past to present. Trop Dis Travel Med Vaccines. 2018;4:14.

57. Rendi-Wagner P, Kollaritsch H. Drug prophylaxis for travelers’ diarrhea. Clin Infect Dis. 2002;34(5):628–33.

58. Duan Q, Huang J, Xiao N, Seo H, Zhang W. Neutralizing Anti-Heat-Stable Toxin (STa) Antibodies Derived from Enterotoxigenic Escherichia coli Toxoid Fusions with STa Proteins Containing N12S, L9A/N12S, or N12S/A14T Mutations Show Little Cross-Reactivity with Guanylin or Uroguanylin. Appl Environ Microbiol. 2018;84(2).

59. Paton AW, Jennings MP, Morona R, Wang H, Focareta A, Roddam LF, et al. Recombinant probiotics for treatment and prevention of enterotoxigenic Escherichia coli diarrhea. Gastroenterology. 2005;128(5):1219–28.

60. Wraith DC, Goldman M, Lambert PH. Vaccination and autoimmune disease: what is the evidence? Lancet. 2003;362(9396):1659–66.

61. Ozen M, Dinleyici EC. The history of probiotics: the untold story. Benef Microbes. 2015;6(2):159–65.

62. Matera M. Bifidobacteria, Lactobacilli… when, how and why to use them. Global Pediatrics. 2024;Volume 8.

63. Celestine Sau-Chan Tham K-KP, Rajeev Bhat & Min-Tze Liong. Probiotic properties of bifidobacteria and lactobacilli isolated from local dairy products. Annals of Microbiology. 22 September 2011;62(September 2012):1079–87.

64. Alok A, Singh ID, Singh S, Kishore M, Jha PC, Iqubal MA. Probiotics: A New Era of Biotherapy. Adv Biomed Res. 2017;6:31.

65. Brubaker J, Zhang X, Bourgeois AL, Harro C, Sack DA, Chakraborty S. Intestinal and systemic inflammation induced by symptomatic and asymptomatic enterotoxigenic E. coli infection and impact on intestinal colonization and ETEC specific immune responses in an experimental human challenge model. Gut Microbes. 2021;13(1):1–13.

66. Thapa HB, Kohl P, Zingl FG, Fleischhacker D, Wolinski H, Kufer TA, et al. Characterization of the Inflammatory Response Evoked by Bacterial Membrane Vesicles in Intestinal Cells Reveals an RIPK2-Dependent Activation by Enterotoxigenic Escherichia coli Vesicles. Microbiol Spectr. 2023;11(4):e0111523.

67. Asfaha S, Dubeykovskiy AN, Tomita H, Yang X, Stokes S, Shibata W, et al. Mice that express human interleukin-8 have increased mobilization of immature myeloid cells, which exacerbates inflammation and accelerates colon carcinogenesis. Gastroenterology. 2013;144(1):155–66.

68. Li W, Zhang S, Wang Y, Bian H, Yu S, Huang L, et al. Complex probiotics alleviate ampicillin-induced antibiotic-associated diarrhea in mice. Front Microbiol. 2023;14:1156058.

69. Asahara T, Takahashi A, Yuki N, Kaji R, Takahashi T, Nomoto K. Protective Effect of a Synbiotic against Multidrug-Resistant Acinetobacter baumannii in a Murine Infection Model. Antimicrob Agents Chemother. 2016;60(5):3041–50.

70. Xu C, Yan S, Guo Y, Qiao L, Ma L, Dou X, et al. Lactobacillus casei ATCC 393 alleviates Enterotoxigenic Escherichia coli K88-induced intestinal barrier dysfunction via TLRs/mast cells pathway. Life Sci. 2020;244:117281.

71. Ahmadi Rouzbahani H, Mousavi Gargari SL, Nazarian S, Abdollahi S. Protective Immunity Against Enterotoxigenic Escherichia coli by Oral Vaccination of Engineered Lactococcus lactis. Curr Microbiol. 2021;78(9):3464–73.

72. Bokoliya SC, Dorsett Y, Panier H, Zhou Y. Procedures for Fecal Microbiota Transplantation in Murine Microbiome Studies. Front Cell Infect Microbiol. 2021;11:711055.

73. Wang T, Teng K, Liu G, Liu Y, Zhang J, Zhang X, et al. Lactobacillus reuteri HCM2 protects mice against Enterotoxigenic Escherichia coli through modulation of gut microbiota. Sci Rep. 2018;8(1):17485.

74. Chandrasekaran P, Weiskirchen S, Weiskirchen R. Effects of Probiotics on Gut Microbiota: An Overview. Int J Mol Sci. 2024;25(11).

